# Longevity interventions in Titan mice attenuate frailty and senescence accumulation

**DOI:** 10.1101/2023.09.08.556694

**Authors:** Benedikt Gille, Annika Müller-Eigner, Shari Gottschalk, Erika Wytrwat, Martina Langhammer, Shahaf Peleg

## Abstract

Wild-type murine models for aging research have lifespans of several years, which results in long experimental duration and late output. Here we explore the short-lived non-inbred Titan mouse as a mouse model to test longevity interventions. We show that Titan mice exhibit increased frailty and cellular senescence at an early age. Dietary intervention attenuates the frailty progression of Titan mice. Additionally, cyclic administration of the senolytic drug Navitoclax at early age increases the lifespan and reduces cellular senescence. Our data suggests that Titan mice can serve as a cost-effective and timely model for longevity interventions in mammals.

## Main part

Murine models were instrumental in many major discoveries in aging research in the last decades. Work in various mouse strains promoted our understanding of the mechanisms of biological aging on all levels of organismal organization and paved the way for aging interventions that aim to extend health- and lifespan^1^. Mice are often used as a key step to test longevity interventions before human clinical studies.

Although mice have shorter lifespans compared to primates, wild-type mouse strains can nonetheless live for several years. This leads to high maintenance costs and long experiment durations. For example, chronic treatment with an expensive drug may lead to significant costs that could hinder the testing of certain interventions. This could ultimately result in a reduction in the number of drugs tested or prevent the testing of costly medications due to limited funding.

Indeed, there are short-lived murine aging models based on genetic modifications that disrupt pathways to mimic progeroid syndromes and lead to premature aging. While genetically modified (GM) short-lived mouse strains yield rapid results and have a predictable phenotype with less variation, they often do not represent a multifactorial aging process as observed in non-GM mice^1^. Additionally, as many GM mouse strains are based on an inbred strain, this could also negatively affect reproducibility and translatability through genetic homogenization^2^. Lastly, GM mice may experience persistent health issues which might be confounding for studying aging. Collectively, utilizing progeria mice that age and die very rapidly may have a sustainable drawback when determining the suitability of a longevity intervention for human studies.

We have previously characterized the non-inbred Titan mouse strain as giant obese mice with a relatively short lifespan. The median and maximum lifespan of Titan mice are 46 and 88 weeks respectively. We have also shown that the mice are responsive to dietary intervention, resulting in improved health metrics and extended lifespan^3^. However, the question of whether these mice could be used to study aging and longevity interventions remained underexplored. While Titan mice are short-lived, we have not assessed the prevalence of hallmarks of aging.

We first quantified age-depended frailty, using the clinical frailty index (FI) as previously described^4^ at regular intervals from 9 to 59 weeks of age. The short lifespan of Titan mice during the FI assessment was comparable to our previous results with a median of 43 weeks (Figure 1a). Importantly, the FI of Titan mice showed a gradual and steady increase, with age being a fixed effect of significant importance in a linear mixed model. Compared to mice at the age of 11 weeks as a baseline, we observed the first significant increase in FI at the age of 17 weeks (Mann-Whitney test with false discovery rate (FDR), P_adjust.signif._ < 0.05; Figure 1b, Supplementary Table 1).

**Figure 1.**
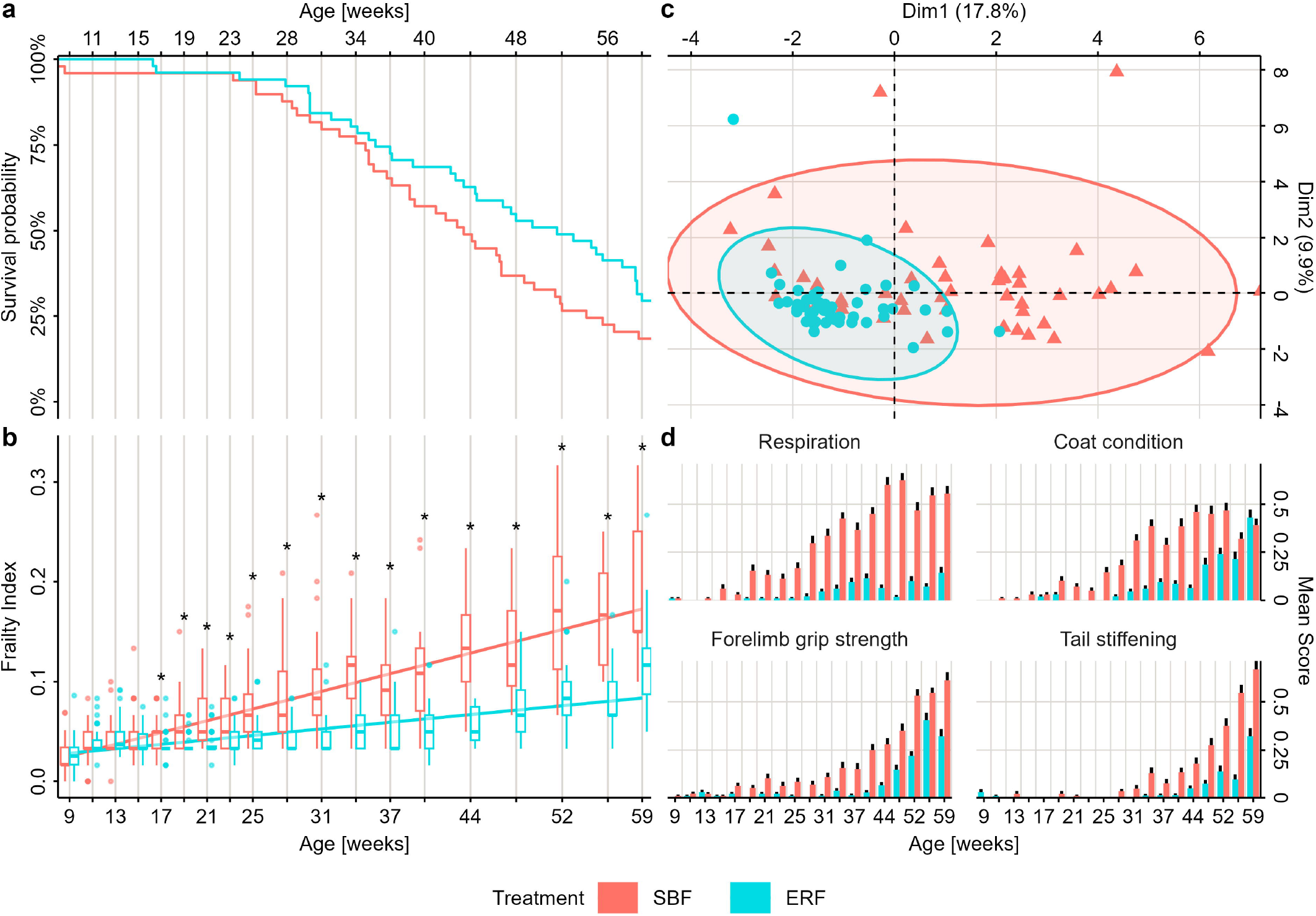
Kaplan-Meier curve and frailty index (FI) assessment of Titan mice under standard breeding feed (SBF) or energy-reduced feed (ERF) regimen. **(a)** Kaplan-Meier curve with survival probability of mice fed with SBF or ERF **(b)** FI boxplots with outliers and linear regression for the diet groups. Asterisks for significance using unpaired Mann-Whitney test and false discovery rate adjustment, P_adjust.signif._ < 0.05. **(c)** Principal component analysis of the FI parameters averages for individual mice. **(d)** Averages and standard error of the top four parameters that caused the clustering of the diet groups.

We have previously shown that dietary intervention of energy-reduced feed (ERF) regime from week 12 increases the lifespan of Titan mice^3^. ERF intervention also significantly reduced the accumulation of deficits expressed in the FI (Figure 1b). Significant differences in FI between diet groups were observable from 17 weeks of age onwards (Mann-Whitney test with FDR, P_adjust.signif._ < 0.05; Figure 1b, Supplementary Table 1). Principal component analysis (PCA) of the FI parameter averages revealed a strong clustering of ERF mice within the broader distribution of SBF mice, predominantly in the dimension of the first principal component. This indicates a stronger uniform frailty phenotype development in ERF-fed mice within the heterogenic frailty characteristics of SBF-fed mice (Figure 1c). Specifically, four FI parameters contributing to the distinction between diet groups were respiration, coat condition, tail stiffening, and forelimb grip strength (Figure 1d). Indeed, loss of tail motility and grip strength indicate early progression of sarcopenia, a typical aging hallmark^5^. Finally, several other FI parameters like gait disorder, hearing loss, and piloerection were increased with aging, while several other parameters, such as cataracts, alopecia, and kyphosis do not appear to correlate with age in these mice (Supplementary Fig. 1). The body weight did not have a significant effect on FI, which indicates that the weight loss effects of ERF did not mediate FI reduction directly (Supplementary Fig. 2a, Supplementary Table 1).

We next characterized the accumulation of senescence, which is a key hallmark of aging. Cellular senescence (CS) is characterized by permanent growth and proliferative arrest, a senescence-associated secretory phenotype^6^, and a loss of regenerative potential^7^. The accumulation of CS during aging, promoted by obesity and accompanied by their systemic effects, is a major contributor to the onset of age-related chronic diseases in mice and humans^8,9^. Indeed, we found that the obese Titan mice show increased senescence-associated beta-galactosidase (SA-β-gal) activity in the liver already at the age of 30 weeks compared to 8-week-old mice. (Student’s t-test, P = 0.036; Fig. 2a and b, Supplementary Table 1).

**Figure 2.**
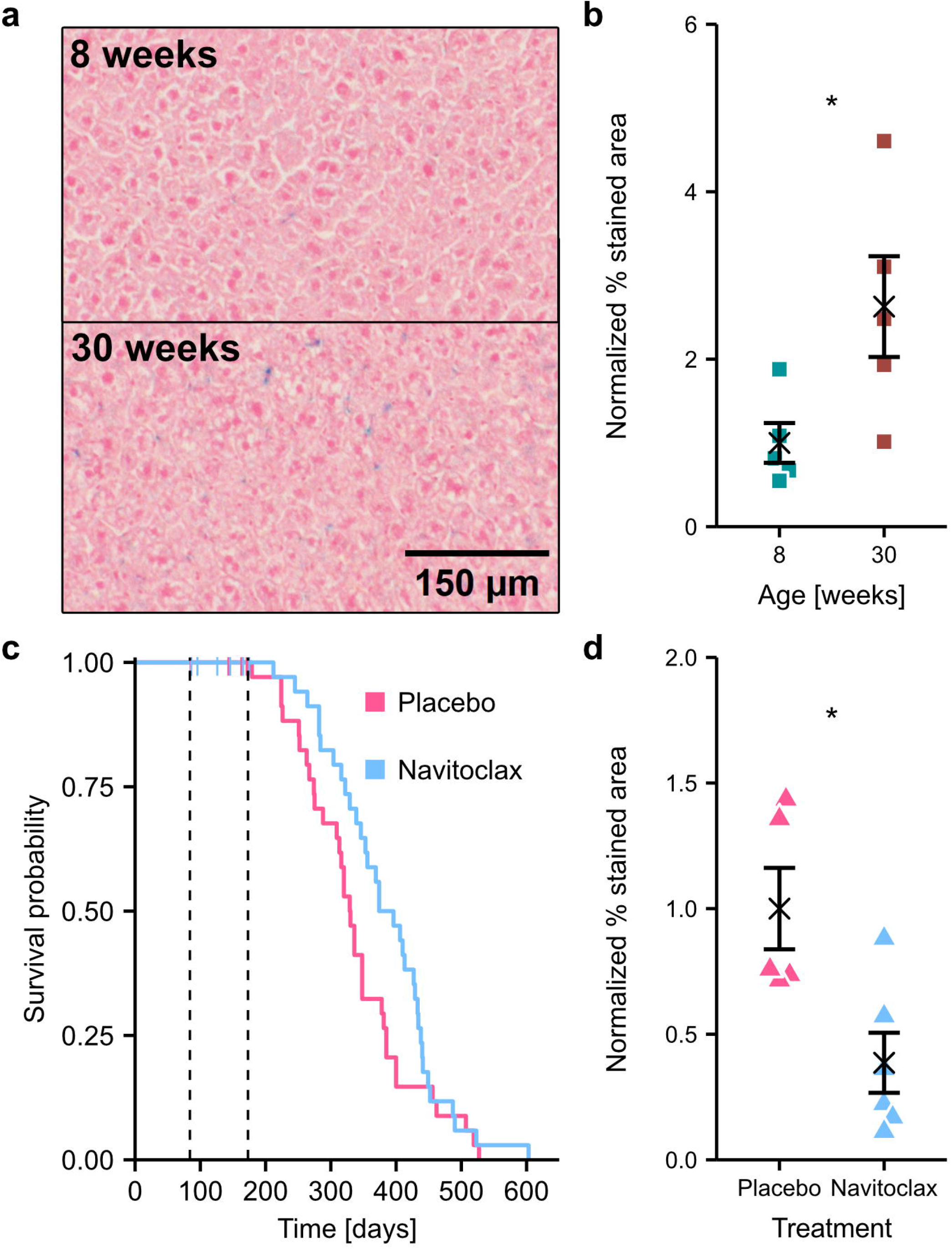
Navitoclax treatment increases lifespan of Titan mice. **(a)** SA-β-gal staining of liver shows increased staining of 8-week-old vs 30-week-old Titan mice. **(b)** Quantification of (a), Student’s t-test, P = 0.036. **(c)** Kaplan-Meier curve of Navitoclax vs placebo. Vertical lines indicate censored mice. Gehan-Breslow-Wilcoxon test, P = 0.026. **(d)** Navitoclax treatment reduced SA-β-gal staining of liver at 26-27 week-old Titan mice, Mann-Whitney test, P = 0.03.

CS is an important target for interventions to slow down or reset the aging processes^8,10^. It forms when damaged cells do not enter apoptosis, but activate cell-cycle arrest pathways and protect themselves from Caspase-9 mediated cell death by an upregulation of the anti-apoptotic BCL-2 family proteins^8,11^. We have therefore tested if senolytic treatment with the BCL-2 inhibitor Navitoclax can extend the lifespan of male Titan mice. Previously, Navitoclax has been shown to induce apoptosis selectively in senescent cells by inhibiting BCL-2 family proteins^12^. Navitoclax is orally bioavailable and well tolerated with minor side effects, like reversible thrombocytopenia^12,13^. Fielder et al. showed that transient treatment with Navitoclax was sufficient to partially ameliorate the detrimental effects of radiation-induced senescence in mice^14^.

We treated 12-week-old Titan mice with Navitoclax for five days, followed by a 16-day break. This was repeated five times to the age of 27 weeks. The treatment resulted in 16.7 % increased median lifespan compared to the placebo group, (330 days placebo vs 385 days Navitoclax) and a significant extension of overall lifespan (Gehan-Breslow-Wilcoxon test, P = 0.026; Fig. 2c). Caspase-9 activity in the liver and blood cell composition, including platelet count, remained unchanged two weeks after the last transient Navitoclax treatment, whereas body weight was reduced by treatment, especially in late life (Supplementary Fig. 2b-h, Supplementary Table 1). Notably, Navitoclax-treated mice show reduced levels of SA-β-gal in the liver compared to the placebo group at the age of 26 to 27 weeks, two weeks after the last treatment (Mann-Whitney test, P = 0.03; Fig. 2d), confirming the results in *C. elegans*^*15*^.

In summary, our data support the notion that male Titan mice display an increased prevalence of hallmarks of aging already at a few months of age. A dietary intervention results in a meaningful attenuation of frailty metrics, thus indicating an improved healthspan in the mice. Additionally, cyclic and transient Navitoclax treatment results in a significant lifespan extension in Titan mice. According to this study, utilizing Titan mice as a model for longevity interventions can expedite translational drug experiments on mammals, which are typically a time-consuming and expensive undertaking.

## Material and Methods

### Animals and husbandry conditions

All experiments and procedures were approved by the state of Mecklenburg-Western Pomerania. The work with mice and health monitoring were conducted in conformity with European FELASA recommendations. For all experiments, mice received water and SBF (V1124-300, ssniff Spezialdi ä ten GmbH) *ad libitum* or, if indicated ERF (V1574-300, ssniff Spezialitäten GmbH)^3^. Housing facilities were maintained at a temperature of 22.5 ± 0.2 °C, at least 50 % humidity, and a 12:12h light-dark cycle. Until the beginning of the experiments, mice were kept in specific-pathogen-free conditions and later transferred to a conventional housing but remained pathogen-free according to FELASA recommendations.

### Frailty Index

For the assessment of the FI, 100 male Titan mice were housed in groups of two in Eurostandard Typ II L cages (Tecniplast, Germany) with nesting material. For less invasive handling and enrichment during FI assessment, acrylic glass tunnels were placed in cages. SBF was given to 49 mice throughout their whole life as a control, while 51 mice received ERF from the age of 12 weeks. Deaths in the Kaplan-Meier function represent both, deceased mice and mice sacrificed due to humane endpoints. FI was evaluated regularly from 9 weeks to 59 weeks, with frequency decreasing over time. Short intervals (every two weeks) were chosen to record changes in early development at higher resolution and later extended (every three, later every four weeks) to reduce handling stress in mice. Based on a procedure by Whitehead et al., a total of 30 health- and aging-associated parameters were scored as 0 (no deficit), 0.5 (mild deficit), or 1 (severe deficit) in every mouse at each time point and the FI was calculated as the mean of the 30 parameter scores^4^.

### SA-β-gal staining

Mouse liver was fixed in phosphate-buffered 10 % formaldehyde at 4 °C for three hours and afterwards cryoprotected by incubation in 30% sucrose in PBS solution at 4 °C for 27 h. For cryosectioning, samples were embedded in OCT and stored at -80 °C for less than 4 weeks. After cutting 8 μm sections at -20 °C in a cryotome, sections were mounted on gelatin-coated slides. Based on manufacturer’s protocol (Senescence β-Galactosidase Staining Kit #9860, Cell Signaling Technology), 200 μl of the supplied fixative solution was added to every section and incubated for 10 min at RT, rinsed with PBS and then covered in 200 μl β-galactosidase staining solution for 15 h at 37 °C. Contrasting counterstaining was done with nuclear fast red for 10 min according to manufacturer’s instructions (Nuclear fast red aluminum sulfate solution, N069.1, Carl Roth), followed by clearing with xylene-substitute (Roticlear, A538.1, Carl Roth) and mounting with toluene-based resin mounting medium (Permount Mounting Medium, Fisher Chemical). Bright-field microscope (Axio Imager A1, Carl Zeiss Microscopy) was used for imaging with 100x magnification and z-stacking, if appropriate. After excluding vessels and artifacts in the images, the percentage of the stained area was measured in three randomized red channel images per individual using a mean of the thresholding algorithms Shanbhag, Intermodes, and MaxEntropy, excluding outlying algorithm results. Analyses were performed in the ImageJ distribution Fiji^16^.

### Navitoclax treatment

To test long-term lifespan effects and senescence marker alterations by senolytic Navitoclax treatment, 40 male Titan mice received 50 mg/kg Navitoclax dissolved in 60 % Phosal50 PG, 30 % polyethyleneglycol and 10 % ethanol through a gavage, while a control group of 40 mice received just the carrier solution as placebo. Mice were housed in groups of two. The treatment phase started at the age of 12 weeks with five days followed by 16 days of break. The drug was administered for five cycles until the mice reached 27 weeks of age. The person administrating the gavage was blinded to the grouping of the mice. Mice that died during the drug administration period were censored in further analysis. SA-β-gal, blood parameters, and Caspase-9 activity were measured in mice 16 days after the last treatment.

### Caspase-9 activity assay

Caspase-9 activity was measured in liver lysate according to manufacturer’s protocol (Merck Caspase-9 Colorimetric Activity Assay Kit APT139, EMD Millipore Corporation, USA). In brief, approximately 50 mg of liver tissue was lysed in provided lysis buffer. After centrifugation, supernatant was mixed with provided assay buffer and Caspase-9 substrate, incubated for 2 h at 37 °C, and measured at 405 nm in a microtiter plate reader.

### Statistics

All statistical analyses were performed in R 4.3.0^17^. The FI data was modeled in a maximum likelihood fitted linear mixed-effect model with lme4 (v1.1-33)^18^, with the complete model consisting of the fixed effects diet group and age, and the random effects weight and individual. To determine the significance of mixed and random effects in a likelihood ratio test, an ANOVA was utilized^19^. Normal distribution was tested with Shapiro-Wilk test^20^. Group-wise comparisons of FI data were performed using the non-parametric Mann-Whitney test with subsequent FDR adjustment^21^. For unsupervised PCA, the individual overall mean scores of every parameter were calculated and then mapped and visualized using the packages FactoMineR (v2.8)^22^and factoextra (v1.0.7)^23^. SA-β-gal data was tested with the non-parametric Mann-Whitney test or Student’s t-test if appropriate^21,24^. Navitoclax treatment was expected to have a stronger impact on early lifespan due to a rebound of progeroid effects of CS after the treatment ended. Therefore, the weighted Gehan-Breslow-Wilcoxon test was used to compare survival functions in the Navitoclax experiment. Survival analysis was performed using the packages survival (v3.5-5)^25^and survminer (v0.4.9)^26^. Blood parameters and Caspase-9 activity data were tested with Student’s t-test. Significance level for all tests was P > 0.05. All descriptive statistical data and test results are summarized in Supplementary Table 1.

## Author Contributions

S.P. designed the research. B.G. performed the experiments and statistical analyses. A.M.G and S.G contributed carrying out the experiments. E.W and M.L supervised the mice work. B.G. and S.P wrote the manuscript with input from the other authors. All authors have read and agreed to the published version of the manuscript.

## Acknowledgments

We would like to thank our technicians Verena Hofer-Pretz and Paula Becker for performing many of the experiments for this study, as well as for managing the laboratory conditions that enabled this work. The authors thank Ines Müntzel, Karin Ullerich, Sabine Maibohm, Benita Lucht, Hildburg Meier, Lisa Rosenow, and Diane Loth from the mouse facility for the animal care and technical assistance.

---

**Supplementary Figure 1.**
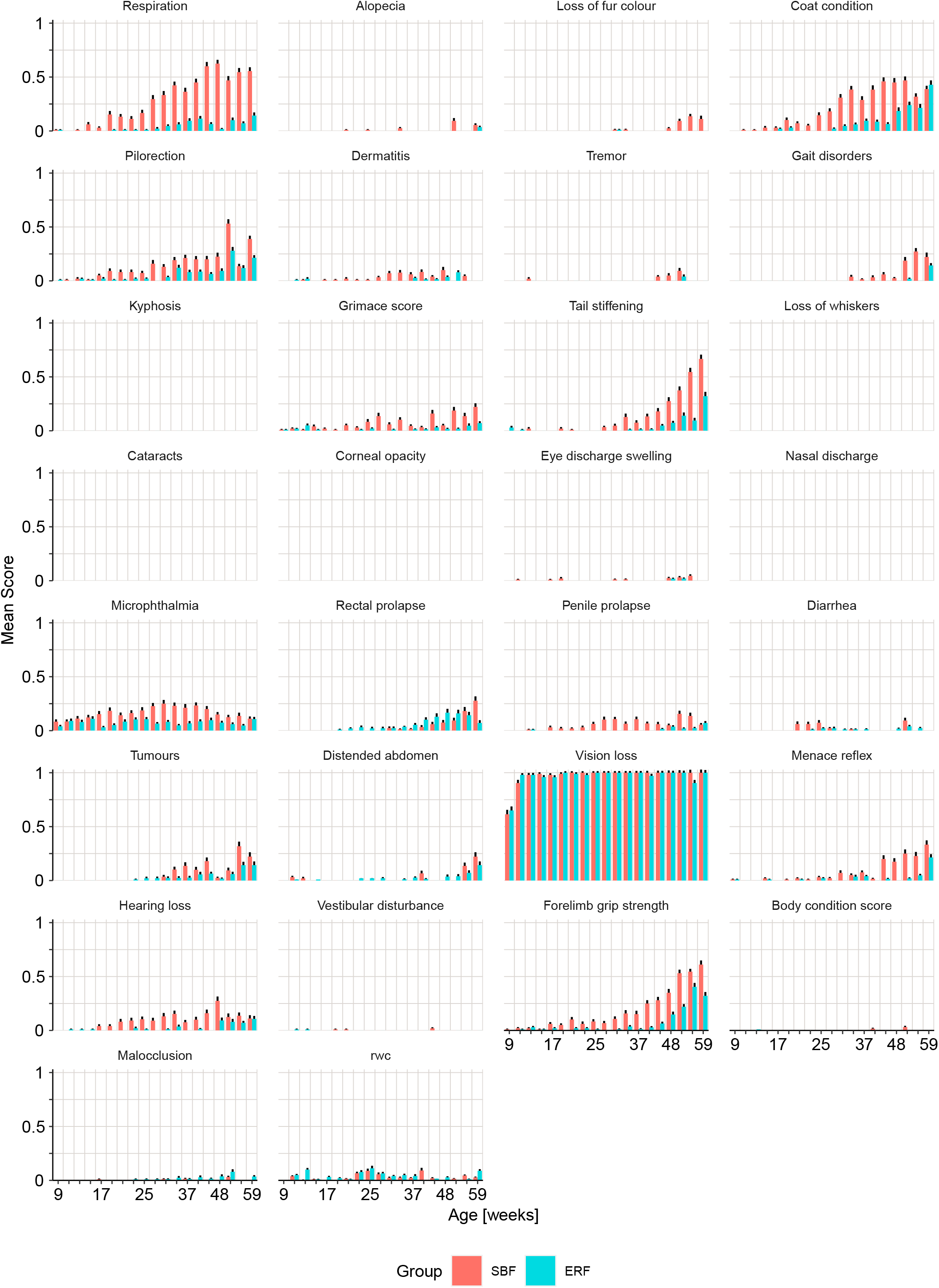
Quantification of individual FI parameters between 9 to 59-week-old Titan mice. While some parameters showed a gradual increase with time, others were not detectable in Titan mice. Bars indicate group mean, error bars represent standard error.

**Supplementary Figure 2.**
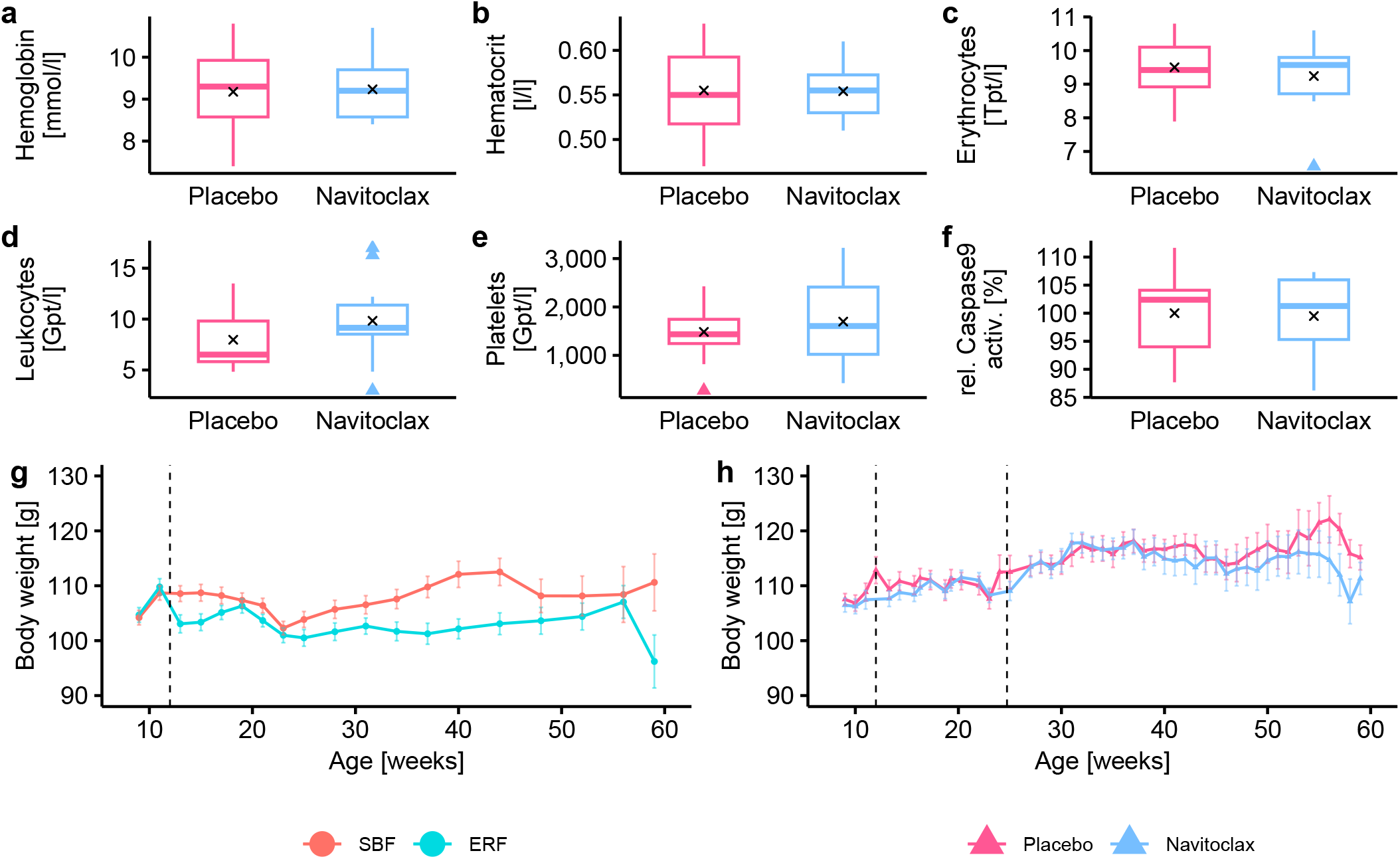
Weight and biochemical parameters. **(a)** Weight of Titan mice during the Frailty Index experiment. Dietary intervention with energy-reduced feed (ERF) reduced the weight of mice significantly compared to standard breeding feed (SBF), linear model, P < 0.001 **(b)** Weight of Titan mice treated with either Navitoclax or placebo. Navitoclax treatment resulted in lower body weight, predominantly in late life, linear model, P = 0.0012 **(c)** Hemoglobin concentration **(d)** Hematocrit proportion **(e)** Erythrocyte count **(f)** Leukocyte count **(g)** Platelet count **(h)** Relative Caspase-9 activity.

