## Supplementary Table 1 for "Longevity interventions in Titan mice attenuate frailty and senescence accumulation"

### Supplementary Table 1 Descriptive statistics and analyses.

Comprehensive list of descriptive measures and statistical tests conducted throughout this study. Standard deviation (SD), standard breed feed (SBF), energy-reduced feed (ERF), not significant (ns), linear mixed model (LMM), false discovery rate (FDR), frailty index (FI), senescence-associated β-galactosidase (SA-β-gal).

| Experiment | Parameter | constants | Factor | Levels | Sample size | Mean ± SD/ median for survival data | Statistical test | P-value/ Adjusted P-value† | Significance summary |
|---|---|---|---|---|---|---|---|---|---|
| Survival curve | Lifespan in days | Males | Diet | SBF | 49 | 304 | log-rank | 0.1006 | ns |
| | | | | ERF | 51 | 361 | | | |
| Frailty Index | FI | Males | Age | - | 100 | - | Likelihood ratio test of LMM | < 0.005 | ** |
| Frailty Index | FI | Males | Diet | SBF | 49 | - | Likelihood ratio test of LMM | 0.013 | * |
| | | | | ERF | 51 | - | | | |
| Frailty Index | FI | Males | Body weight | - | - | - | Likelihood ratio test of LMM | 0.65 | ns |
| Frailty Index | FI | Males | Age [weeks] | 11 | 47 | 0.038 ± 0.017 | Multiple unpaired Mann-Whitney tests with FDR adjustment | †0.2 | ns |
| | | | | 13 | 49 | 0.042 ± 0.017 | | | |
| | | | | 11 | 47 | 0.038 ± 0.017 | | †0.19 | ns |
| | | | | 15 | 49 | 0.043 ± 0.02 | | | |
| | | | | 11 | 47 | 0.038 ± 0.017 | | †0.002 | * |
| | | | | 17 | 49 | 0.048 ± 0.017 | | | |
| | | | | 11 | 47 | 0.038 ± 0.017 | | < †0.001 | ** |
| | | | | 19 | 49 | 0.057 ± 0.025 | | | |
| | | | | 11 | 47 | 0.038 ± 0.017 | | < †0.001 | ** |
| | | | | 21 | 49 | 0.061 ± 0.026 | | | |
| | | | | 11 | 47 | 0.038 ± 0.017 | | < †0.001 | ** |
| | | | | 23 | 49 | 0.059 ± 0.025 | | | |
| | | | | 11 | 47 | 0.038 ± 0.017 | | < †0.001 | ** |
| | | | | 25 | 48 | 0.071 ± 0.034 | | | |
| | | | | 11 | 47 | 0.038 ± 0.017 | | < †0.001 | ** |
| | | | | 28 | 44 | 0.081 ± 0.041 | | | |
| | | | | 11 | 47 | 0.038 ± 0.017 | | < †0.001 | ** |
| | | | | 31 | 42 | 0.091 ± 0.049 | | | |

| Experiment | Parameter | constants | Factor | Levels | Sample size | Mean ± SD/ median for survival data | Statistical test | P-value/ Adjusted P-value† | Significance summary |
|---|---|---|---|---|---|---|---|---|---|
| | | | | 11 | 47 | 0.038 ± 0.017 | | < †0.001 | ** |
| | | | | 34 | 39 | 0.107 ± 0.039 | | | |
| | | | | 11 | 47 | 0.038 ± 0.017 | | < †0.001 | ** |
| | | | | 37 | 33 | 0.096 ± 0.042 | | | |
| | | | | 11 | 47 | 0.038 ± 0.017 | | < †0.001 | ** |
| | | | | 40 | 30 | 0.11 ± 0.049 | | | |
| | | | | 11 | 47 | 0.038 ± 0.017 | | < †0.001 | ** |
| | | | | 44 | 25 | 0.131 ± 0.049 | | | |
| | | | | 11 | 47 | 0.038 ± 0.017 | | < †0.001 | ** |
| | | | | 48 | 20 | 0.132 ± 0.049 | | | |
| | | | | 11 | 47 | 0.038 ± 0.017 | | †0.002 | * |
| | | | | 52 | 16 | 0.169 ± 0.078 | | | |
| | | | | 11 | 47 | 0.038 ± 0.017 | | †0.007 | * |
| | | | | 56 | 11 | 0.168 ± 0.058 | | | |
| | | | | 11 | 47 | 0.038 ± 0.017 | | †0.016 | * |
| | | | | 59 | 9 | 0.186 ± 0.077 | | | |
| Frailty Index | body weight | Males | Diet | SBF | 49 | - | Linear regression | < 0.001 | ** |
| | | | | ERF | 51 | | | | |
| Frailty Index | FI | Males, 9 weeks old | Diet | SBF | 48 | 0.025 ± 0.019 | Multiple unpaired Mann-Whitney tests with FDR adjustment | †0.912 | ns |
| | | | | ERF | 50 | 0.026 ± 0.018 | | | |
| | | Males, 11 weeks old | | SBF | 47 | 0.038 ± 0.017 | | †0.96 | ns |
| | | | | ERF | 50 | 0.040 ± 0.013 | | | |
| | | Males, 13 weeks old | | SBF | 49 | 0.042 ± 0.017 | | †0.37 | ns |
| | | | | ERF | 50 | 0.044 ± 0.016 | | | |
| | | Males, 15 weeks old | | SBF | 49 | 0.043 ± 0.02 | | †0.42 | ns |
| | | | | ERF | 50 | 0.038 ± 0.011 | | | |
| | | Males, 17 weeks old | | SBF | 49 | 0.048 ± 0.017 | | < †0.001 | ** |
| | | | | ERF | 48 | 0.036 ± 0.011 | | | |
| | | Males, 19 weeks old | | SBF | 49 | 0.057 ± 0.025 | | < †0.001 | ** |
| | | | | ERF | 48 | 0.038 ± 0.009 | | | |
| | | | | SBF | 49 | 0.061 ± 0.026 | | < †0.001 | ** |

| Experiment | Parameter | constants | Factor | Levels | Sample size | Mean ± SD/ median for survival data | Statistical test | P-value/ Adjusted P-value† | Significance summary |
|---|---|---|---|---|---|---|---|---|---|
| | | Males, 21 weeks old | | ERF | 48 | 0.038 ± 0.011 | | | |
| | | Males, 23 weeks old | | SBF | 49 | 0.059 ± 0.025 | | †0.004 | * |
| | | | | ERF | 48 | 0.044 ± 0.015 | | | |
| | | Males, 25 weeks old | | SBF | 48 | 0.071 ± 0.034 | | < †0.001 | ** |
| | | | | ERF | 46 | 0.047 ± 0.018 | | | |
| | | Males, 28 weeks old | | SBF | 44 | 0.081 ± 0.041 | | < †0.001 | ** |
| | | | | ERF | 46 | 0.042 ± 0.014 | | | |
| | | Males, 31 weeks old | | SBF | 42 | 0.091 ± 0.049 | | < †0.001 | ** |
| | | | | ERF | 43 | 0.046 ± 0.019 | | | |
| | | Males, 34 weeks old | | SBF | 39 | 0.107 ± 0.039 | | < †0.001 | ** |
| | | | | ERF | 41 | 0.052 ± 0.022 | | | |
| | | Males, 37 weeks old | | SBF | 33 | 0.096 ± 0.042 | | < †0.001 | ** |
| | | | | ERF | 37 | 0.054 ± 0.027 | | | |
| | | Males, 40 weeks old | | SBF | 30 | 0.11 ± 0.049 | | < †0.001 | ** |
| | | | | ERF | 35 | 0.054 ± 0.023 | | | |
| | | Males, 44 weeks old | | SBF | 25 | 0.131 ± 0.049 | | < †0.001 | ** |
| | | | | ERF | 31 | 0.057 ± 0.019 | | | |
| | | Males, 48 weeks old | | SBF | 20 | 0.132 ± 0.049 | | < †0.001 | ** |
| | | | | ERF | 27 | 0.07 ± 0.033 | | | |
| | | Males, 52 weeks old | | SBF | 16 | 0.169 ± 0.078 | | < †0.001 | ** |
| | | | | ERF | 25 | 0.091 ± 0.04 | | | |
| | | Males, 56 weeks old | | SBF | 11 | 0.168 ± 0.058 | | < †0.001 | ** |
| | | | | ERF | 21 | 0.081 ± 0.034 | | | |
| | | Males, 59 weeks old | | SBF | 9 | 0.186 ± 0.077 | | †0.041 | * |
| | | | | ERF | 14 | 0.122 ± 0.061 | | | |
| SA-β-gal | % stained area | Males, SBF | age in weeks | 8 | 5 | 0.034 ± 0.018 | Student's t-test | 0.036 | * |
| | | | | 30 | 5 | 0.09 ± 0.046 | | | |
| | | Males | Treatment | Placebo | 5 | 2.19 ± 0.794 | two-sided Mann-Whitney test | 0.03 | * |
| | | | | Navitoclax | 6 | 0.846 ± 0.641 | | | |
| Navitoclax | lifespan in days | Males | Treatment | Placebo | 40 | 330 | | 0.026 | * |

| Experiment | Parameter | constants | Factor | Levels | Sample size | Mean ± SD/ median for survival data | Statistical test | P-value/ Adjusted P-value† | Significance summary |
|---|---|---|---|---|---|---|---|---|---|
| | | | | Verum | 40 | 385 | Gehan-Breslow-Wilcoxon | | |
| Navitoclax | body weight | Males | Treatment | Placebo | 40 | - | Linear regression | 0.0012 | * |
| | | | | Verum | 40 | | | | |
| Navitoclax | Hemoglobin | Males | Treatment | Placebo | 12 | 9.175 ± 1.048 | Student's t-test | †0.877 | ns |
| | | | | Verum | 12 | 9.233 ± 0.751 | | | |
| Navitoclax | Hematocrit | Males | Treatment | Placebo | 12 | 0.555 ± 0.052 | | †0.962 | ns |
| | | | | Verum | 12 | 0.554 ± 0.032 | | | |
| Navitoclax | Erythrocytes | Males | Treatment | Placebo | 12 | 9.502 ± 0.929 | | †0.529 | ns |
| | | | | Verum | 12 | 9.242 ± 1.06 | | | |
| Navitoclax | Leukocytes | Males | Treatment | Placebo | 12 | 7.97 ± 2.861 | | †0.205 | ns |
| | | | | Verum | 12 | 9.836 ± 4.044 | | | |
| Navitoclax | Platelets | Males | Treatment | Placebo | 12 | 1,484.5 ± 619.018 | | †0.514 | ns |
| | | | | Verum | 12 | 1,697.25 ± 922.22 | | | |
| Navitoclax | rel. Caspase-9 activity | Males | Treatment | Placebo | 6 | 100 ± 8.848 | | †0.919 | ns |
| | | | | Verum | 6 | 99.49 ± 8.202 | | | |
